# Evaluating the Influence of Disease-Gene Associations in the Significance of Disease Modules through the lens of Network Medicine

**DOI:** 10.1101/2025.06.05.657982

**Authors:** Antonio Gil Hoed, Lucía Prieto-Santamaría, Alejandro Rodríguez-González

## Abstract

The rapid expansion of genomic and biomedical data has paved the way for constructing complex disease networks, offering new insights into disease mechanisms and therapeutic target identification. Central to these networks are disease modules, which are constructed from seed genes that are prioritized based on their relevance to a given disease. Gene prioritization aims to rank genes based on their strength of association with a disease. Resources like DisGeNET, integrated within DISNET knowledge base, assigns a Gene-Disease Association (GDA) score that reflects the confidence in the association between a gene and a disease. This study investigates how both disease module sizes and GDA scores influence the statistical significance of these modules. By characterizing disease modules, filtering the data based on their GDA score thresholds, and analysing the relationship between their size, GDA score, and significance, potential cutoffs for robust module construction are estimated. Our findings suggest that disease modules filtered to include GDA scores above 0.3 and module sizes greater than 10 tend to be significant. These insights provide guidance for optimal gene prioritization and module selection, ultimately enhancing strategies for target identification and drug repurposing in network medicine.

## I. Introduction

Complex biological systems can be modelled as networks or graphs, where nodes represent genes or proteins, and edges depict their physical or functional relationships. This network-based approaches present relevant clinical applications, such us disease classification, target prioritization or drug repurposing [1].

A disease module is a connected sub-network within a biological network, such as the interactome, where gene products (e.g., proteins), represented as nodes, associated with the same phenotype tend to interact and cluster together. Proper identification of disease modules helps uncover new disease-related genes and pathways, facilitating identification of drug targets [2].

To accurately identify and construct a disease module, it is essential to determine which genes function as seed disease genes and contribute to the observed phenotype. Gene prioritization plays a crucial role in identifying causative genes, enabling the selection of the most relevant candidates for further analysis [3].

Advances in genomics have produced vast amounts of data employed to augment the interactome, and the number of disease seed genes. However, this abundance also presents challenges, such as the integration of heterogeneous data from multiple sources and the extraction of the most critical information [4]. To determine which genes have a stronger association with a specific phenotype, some databases such us DisGeNET, incorporate a Gene-Disease Association (GDA) score, which symbolizes the degree of connection between a gene and a disease, assigning higher values to more robust and curated associations [5].

Since disease-associated proteins tend to interact, alternative methods for identifying disease-related genes rely on analyzing their relationships with neighboring proteins to determine which genes are truly involved in the disease phenotype [6]. Many disease module algorithms leverage this principle, incorporating interactome proteins into the disease module based on their network connections [7].

The main objective of this research is to identify which seed disease genes should be included, and which parameters contribute to the significance of the disease module. This is achieved by determining several factors: the GDA score threshold, filtering disease seed genes with this approach, and the size of the resulting disease module (that is, the number of proteins in it). **Fig. 1** includes the graphical abstract of the methodology employed in this study.

**Fig. 1.**
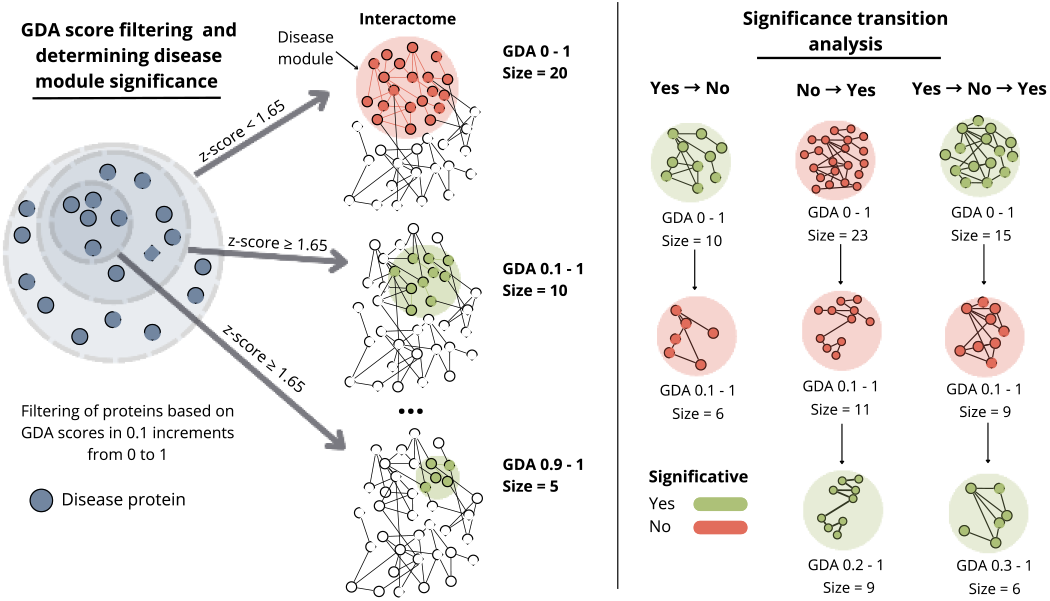
Summary of the analysis conducted in this paper. 1) Filtering of disease-associated genes based on their Gene-Disease Association (GDA) score. 2) Extraction of the disease module from the interactome. 3) Evaluation of disease module significance based on statistical thresholds. 4) Transition analysis showing how changes in the GDA score and disease module size impact disease module significance.

This paper is structured as follows: Section II reviews the relevant literature in the field of network medicine and disease modules’ characterization. Section III describes the biomedical data and methodology used in this study. Section IV presents and discusses the results. Finally, section V outlines the conclusions drawn from the study’s findings and section VI presents the future lines.

### II. Related works

Network science has evolved from traditional graph theory into a comprehensive framework for understanding the complex, interconnected structures found in diverse systems, going from technological networks to the intricate web of interactions within biological cells [8]. This graph-based approach is also used to analyze complex systems such as the full spectrum of known associations between disease phenotypes and their related genes [9]. Network medicine applies network science to explore the molecular underpinnings of human diseases. By constructing interactomes and identifying disease modules, this approach reveals how perturbations in biological networks contribute to disease pathogenesis [10].

Rather than being caused by a single gene, many diseases result from a cascade of interactions within the biological network, or interactome. Advances in network medicine have been propelled due to the vast availability of high-throughput data and the growing recognition that disease-associated proteins tend to interact within each other, creating distinct “*disease modules*” [11]. Furthermore, network-based analyses of omics data further enhance our understanding by revealing the molecular mechanisms underlying diseases and the patterns of protein interactions [12]. When combined with functional and clinical studies, this approach enhances our understanding of biological functions and disease pathogenesis [2], aiding in prioritizing diagnostic markers and therapeutic candidate genes [13]. Consequently, providing possible drug repurposing scenarios, offering a faster and more cost-effective alternative to traditional drug development [14]. Previous results indicate that diseases involved in successful drug repurposing cases tend to have higher GDA scores with the drug target gene compared to the average in DISNET. This suggests that when both the original and new indications share a strong genetic link with a drug’s target, the biological connection is robust, making the disease a promising candidate for repurposing [15].

Several algorithms have been developed to optimize the identification of this disease modules [11], [16], [17], [18], [19]. One of the first, simpler and more restrictive approaches consist in identifying the Largest Connected Component (LCC), which defines the disease module as the largest group of interconnected disease-associated nodes [12].

Gene prioritization involves assigning similarity or confidence scores to genes and ranking them based on the likelihood of their association with a target disease. This approach is essential for identifying high-confidence candidate genes that may contribute to disease phenotypes. Gene prioritization uses existing biological knowledge—such as gene annotations detailing gene products and their functional and structural properties—to accomplish the task [4]. Disease gene prioritization also leverages the Protein-Protein Interaction (PPI) network and the “guilt-by-association” principle (genes related to disease seed genes tend to be in the disease module) to systematically identify additional candidate genes, making it a key tool in network medicine [6], [20], [21], [22], [23], [24].

Although vast datasets offer valuable insights, they often rely on diverse vocabularies that vary in granularity and origin. As a result, these discordant annotations may inadequately capture the underlying molecular mechanisms, often reflecting historical or symptomatic descriptions rather than true mechanistic details, potentially leading to unreliable conclusions in studies of disease mechanisms. Bird’s-eye-view network medicine approaches leverage large-scale disease association data to uncover global patterns in disease relationships and underlying mechanisms. However, their findings should be complemented with detailed molecular data from well-defined patient cohorts to mitigate biases from phenotype-based definitions. Ultimately, choosing which data to use is crucial, as not all available data can be trusted [25].

## III. Materials and methods

This section includes the data used as the foundation of the study, followed by a detailed explanation of the methodology applied to propose a potential GDA score threshold, minimum number of seed nodes and disease module size to establish a significant disease module.

### A. Materials

The data utilized was acquired from the database of the DISNET ^1^ project, which had the purpose of extracting, integrating, and analyzing biomedical information on diseases (including phenotypic, biologic and pharmacologic data) from public sources. Its main objective was to create a structured and updatable disease network [26]. Among other, DISNET integrated data from DisGeNET, a platform designed to gather and standardize information on disease-associated genes and variants from multiple sources [5], [27]. The DisGeNET score reflects the strength of GDAs by assigning higher values to relationships reported by multiple databases, particularly those curated by experts, and to associations supported by a large body of scientific literature [5]. The PPI network was obtained from neXtProt, a knowledge platform which includes the latest understanding of human proteins [28].

For this study, we used 439,864 protein-protein interactions, 16,460 gene-protein associations and 1,045,745 gene-disease associations with a total of 21,775 diseases. All the files containing these data, as well as the code developed for the next subsection analyses, are accessible through the following GitLab repository^2^.

### B. Methods

The research for optimal GDA score and module size was initiated with the identification of disease modules. To carry out this characterization, the PPI network was generated beforehand. Next, the significance of the obtained disease modules was analyzed, observing how the disease module size and GDA score influenced such significance of the module. Finally, the transitions of significance that disease modules underwent as a result of the GDA score filtering were classified and studied.

#### 1) Interactome

The PPI network, also called the interactome, was constructed using data on protein interactions. In this network, nodes represent proteins, and edges connect proteins that physically or functionally interact with one another.

#### 2) Gene-Disease Association score range

For each disease, all associated genes and their corresponding GDA scores were collected and divided into ranges from 0 to 1, with increments of 0.1 per range. That is, we defined lists of associated genes with their GDA score matching the following ranges: 0–1, 0.1–1, 0.2–1, 0.3–1, 0.4– 1, 0.5–1, 0.6–1, 0.7–1, 0.8–1 and 0.9–1. Such 0.1 increment in the ranges was based on the observed GDA score distribution. This stratification allowed us to examine how module size and significance change with varying GDA score thresholds, progressively filtering out weaker and less curated associations, thus assessing its impact on disease module composition.

#### 3) Disease module obtention

Each disease module was characterized following the methodology outlined by Aldana et al. in 2024 [29], consisting of the next steps:

##### 3.1) Identification of the disease module

For each disease and range, with their corresponding nodes, the LCC was identified in the interactome, representing the highest number of directly interconnected disease proteins. This subgraph is referred to as the disease module, where only disease modules with a size of 5 or above have been considered. Modules consisting in less than 5 proteins do not capture enough biological information.

##### 3.2) Statistical validation of the disease module

To determine the statistical significance of the disease module, a null model with logarithmic binning was used, providing a p-value and z-score for further analysis. This approach, commonly employed in network medicine frameworks, generates random samples as benchmarks, enabling the comparison of observed network measures against these null hypotheses to assess their non-randomness.

##### 3.3) Statistical significance

A disease module was considered statistically significant if it met the criteria of z-score ≥ 1.65. According to Menche et al., 2015, when the z-score exceedes 1.65, the size of the disease module is considered significantly larger than expected by chance [12].

#### 4) Disease module significance based on GDA score range

##### 4.1) Overall module size

We depicted the distributions of all obtained disease modules, considering their GDA score, disease module size, and number of seed genes. A Mann-Whitney U test [30] was performed comparing significant and non-significant values.

##### 4.2) Significance evolution

For each disease, the disease module varies depending on the GDA score range, which can cause the significance of the module to fluctuate, transitioning from significant to non-significant and vice versa. All of the diseases were classified according to the transitions they underwent, and the relationship between disease module size and GDA score range was analyzed.

## IV. Results and discussion

### 1) Disease dataset analysis

The analysis of the 21,775 original diseases and their associated gene counts revealed that most diseases have a low number of associated genes, following the expected power-law distribution, as shown in **Fig. 2**. This observation indicates that a substantial number of diseases will not have enough number of genes to present a significant disease module.

**Fig. 2.**
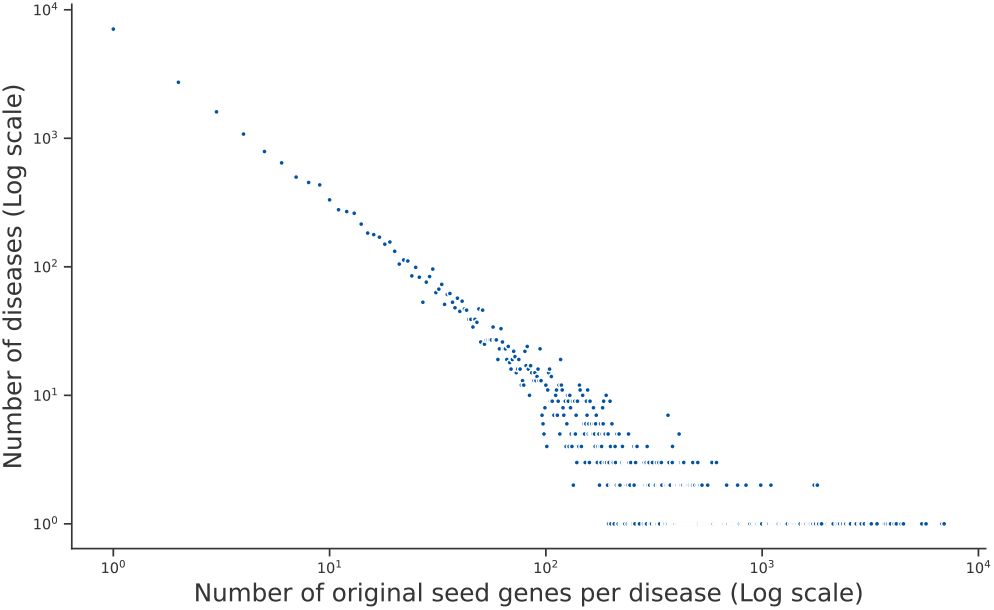
Distribution of the number of seed genes per disease. Scatter plot representing the distribution of diseases according to the number of original seed genes.

GDAs in DisGeNET were compiled from multiple sources and classified into four categories: Literature, Inferred, Animal and Curated. The GDA score derives from an in-house formula that compiles the provenance of each association (it is detailed in the previously linked repository). Literature provided the highest number of associations; however, its reliability is lower compared to other sources. This is reflected in its lower mean GDA score, followed by Inferred and Animal sources. In contrast, Curated sources exhibited the highest GDA score values. A key objective of this study was to assess whether lower GDA scores should be excluded, which would imply considering literature-based data less significant than other sources. As shown in **Fig. 3**, most of the data had a GDA score below 0.3, a factor that may difficult the obtention of a significant disease module.

**Fig. 3.**
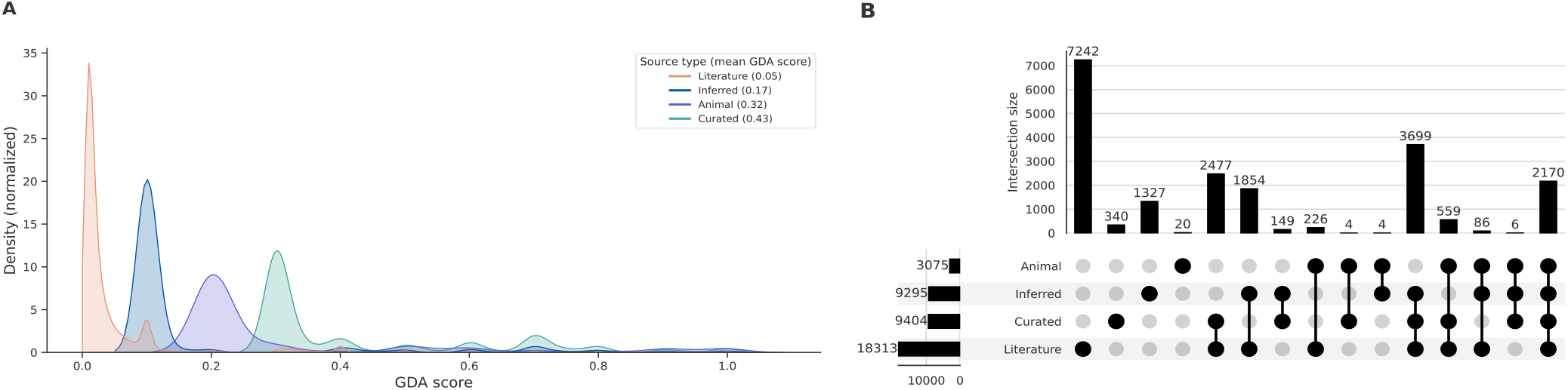
A)Density distribution of Gene-Disease Association (GDA) scores by source type. Density plot displaying the distribution of GDA scores for each source type, each density is independently normalized. The mean GDA score of each plot is indicated in the legend. **B) Gene intersections by source type**. UpSet plot illustrating the intersections of unique GDAs identified for each source type.

### 2) Obtained disease modules analysis

Out of the 21,775 diseases analyzed, only 3,161 had a module size of 5 proteins or more. Among these, without filtering their GDA score, 670 were classified as non-significant, while 2,554 were significant, highlighting a clear disparity between the two groups. This finding underscores the importance of filtering disease modules by size. Without applying the minimum module size threshold of 5, only 5,235 disease modules would have been classified as significant, compared to 16,340 non-significant disease modules.

To examine whether significant and non-significant disease modules differ, a Mann-Whitney U test was performed on three variables: GDA score, number of initial seed genes, and disease module size. The results, depicted in **Fig. 4**, revealed significant differences for all three variables.

**Fig. 4.**
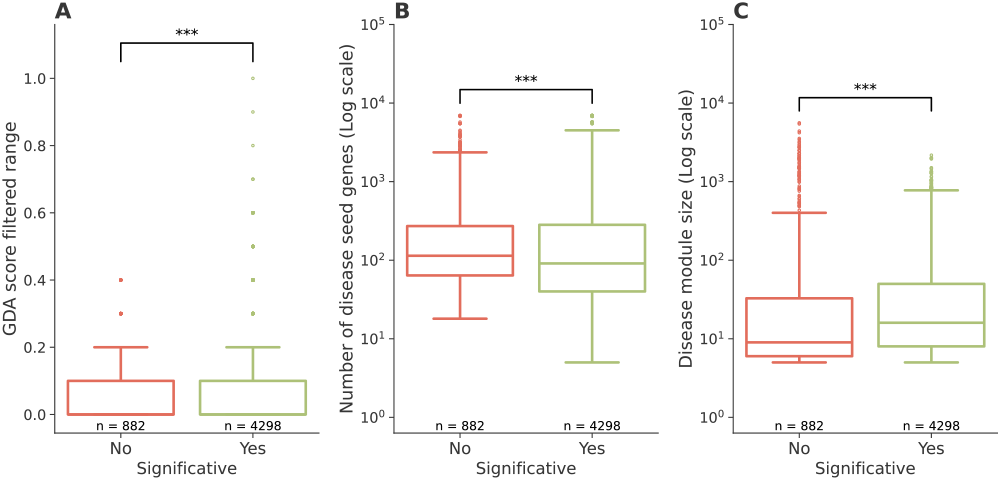
Comparison between significant and non-significant disease modules. Boxplots illustrate the distribution of: (A) Gene-Disease Association (GDA) score range; (B) the number of seed genes; and (C) disease module size, grouped by statistical significance (Yes/No). Each plot shows the median and interquartile range for both categories, with the number of observations (n) indicated below each category. Significance between groups was assessed using the Mann-Whitney U test, and results are annotated with asterisks (*** for p < 0.001, ** for p < 0.01, * for p < 0.05)

For the GDA score ranges, the overall difference appears minimal, as most data points for both significant and non-significant modules fall below a 0.3 value. However, beyond this threshold, a noticeably higher proportion of significant modules emerges. Similarly, the number of seed genes does not exhibit a strong distinction between the two groups. In contrast, disease module size shows a more recognizable difference, with significant disease modules tending to be slightly larger. This suggests that disease module size may play a more critical role than the number of seed genes. While a higher number of initial seed genes is generally associated with a larger disease module, it is not necessarily the determining factor. Moreover, since higher GDA score thresholds favor significant modules, the filtering process naturally results in fewer seed genes. For this analysis, the data was examined per GDA filtered range, meaning a disease could appear multiple times if its disease module remained larger than 5.

Analyzing disease module size across different GDA filtered score ranges, as noticed in **Fig. 5**, for at least half of the values in each category, significant disease modules consistently have a larger size than non-significant ones. However, at a GDA score range of 0.1 (i.e. including genes with 0.1–1 scores), the difference is minimal. From 0.2 onward, the distinction becomes more pronounced. Additionally, the distribution of values across GDA scores reveals an uneven pattern. As previously mentioned, there are more significant values than non-significant ones, but their distribution is not uniform. Proportionally, significant values become more prevalent at higher GDA scores, whereas non-significant values are more concentrated around a GDA score of 0. In fact, no non-significant values are observed above 0.5.

**Fig. 5.**
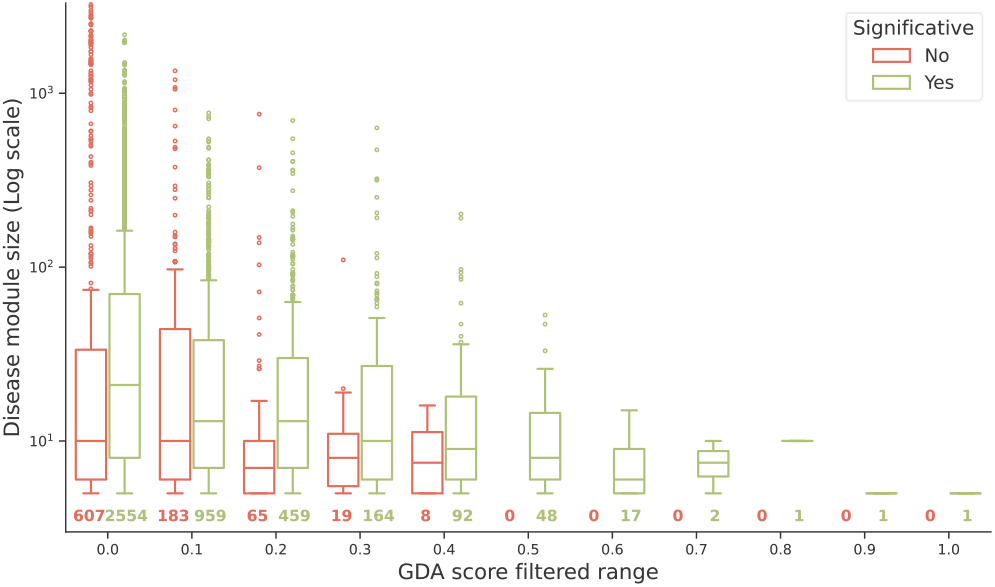
Relationship between Gene-Disease Association (GDA) score range and disease module size. Boxplot illustrating the distribution of disease module sizes across different GDA score ranges (that is, filtering out those below the threshold). The data is categorized based on statistical significance (Yes/No), with the number of diseases in each group annotated below the corresponding boxplots.

This indicates that the higher the GDA score, the less likely it is to find a non-significant disease module. This suggests that sources such as animal models or curated databases are more reliable for constructing disease modules than literature-based sources. This does not mean that literature-based sources are unreliable, but it highlights the need for caution when relying solely on them. Additionally, if a disease module includes any other source type, it is almost certain that it will also incorporate literature-based data.

### 3) Significance transitions based on the GDA score

For each transition type listed in **Table I**, an analysis was conducted (**Fig. 6**). The largest group consisted of consistently significant values, which initially exhibited a high disease module size. As the GDA score increased, the module size gradually decreased due to the reduced availability of seed genes. However, the higher GDA scores appeared to compensate for this decrease, maintaining the overall significance of the disease module.

**TABLE 1.** Observed Significance Transitions IN Disease Modules AND Corresponding Disease Counts.

| Significance transition type | Number of diseases |
| --- | --- |
| Yes | 2463 |
| No | 499 |
| No → Yes | 101 |
| Yes → No | 72 |
| Yes → No → Yes | 17 |
| No → Yes → No | 4 |
| No → Yes → No → Yes | 3 |
| Yes → No → Yes → No | 2 |

**Fig. 6.**
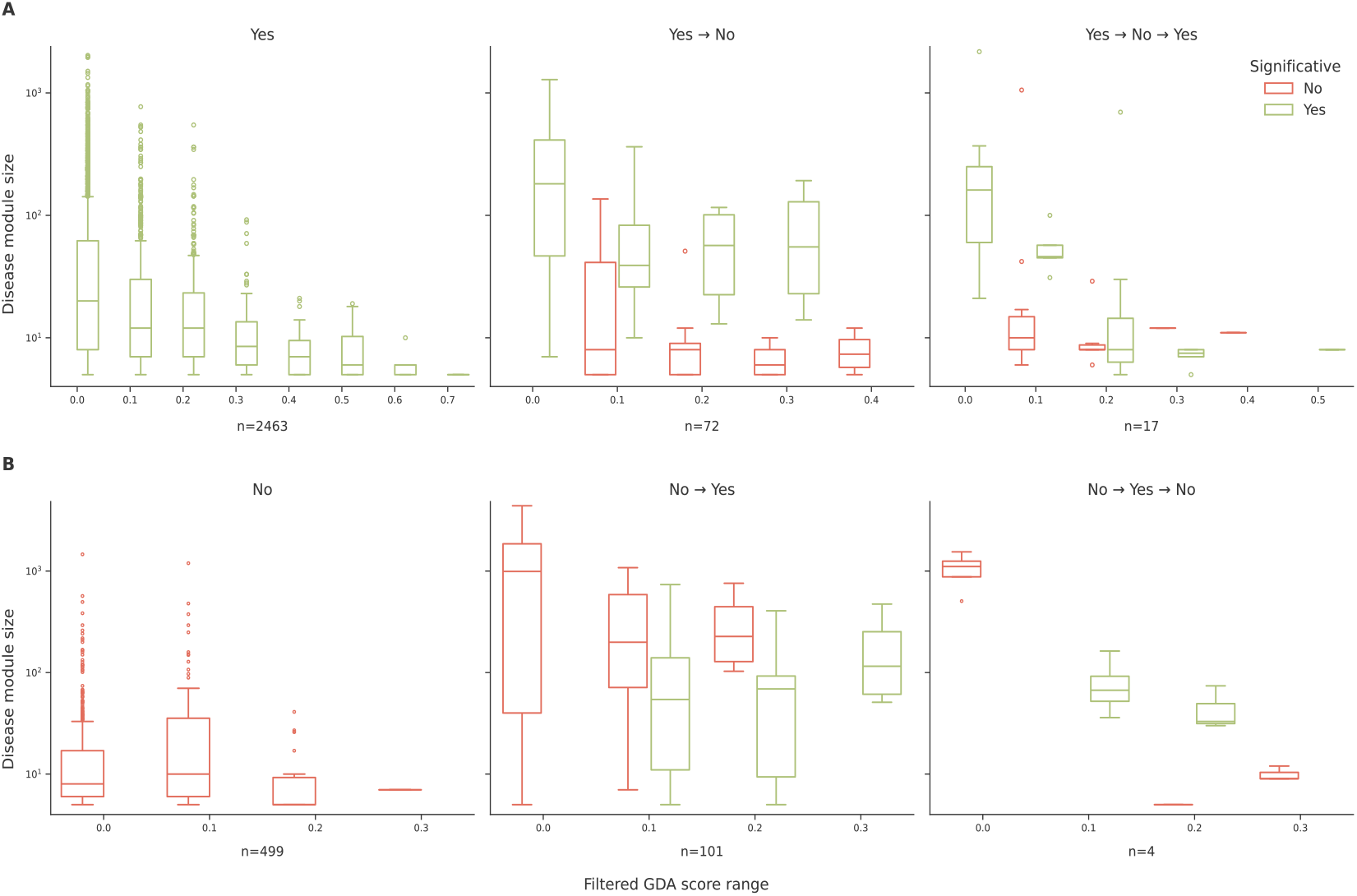
A) Disease module size for transitions beginning with significant values. Boxplots of disease module size (Log scale) by filtered Gene-Disease Association (GDA) score range for “Yes” transitions. Panels show “Yes”, “Yes → No”, and “Yes → No → Yes” scenarios with the number of diseases (n) indicated below. **B) Disease module size for transitions beginning with non-significant values**. Boxplots of disease module size (Log scale) by filtered GDA score range for “No” transitions. Panels represent “No”, “No → Yes”, and “No → Yes → No” scenarios, with n diseases shown below.

Continuing with transitions that begin with significant values, the “Yes → No” transition reveals that significant modules initially exhibit considerably higher values than their non-significant counterparts. This suggests that the loss of significance is primarily driven by a sharp decline in disease module size.

A similar pattern is observed in the “Yes → No → Yes” transition. The initial drop in significance appears to be linked to the decrease in module size. However, upon regaining significance, the disease module does not increase in size, as no additional seed genes are incorporated. In fact, the number of seed genes is reduced. Notably, this transition occurs around GDA thresholds of 0.2 and 0.3, suggesting that filtering genes at these levels enhances significance by eliminating weaker disease-gene associations. This aligns with previous studies, where some authors directly filtered associations below a GDA score of 0.3 [20], [21], given that curated sources are included at this threshold.

For the always non-significant group, the median disease module size remains below 10 across all GDA score values, and the 0.3 GDA score is not exceeded. In contrast, the always significant group surpasses both the disease module size of 10 and the GDA score threshold. This reinforces the idea that disease module size and GDA score plays a crucial role in determining significance.

The “No → Yes” transition presents an interesting case, as most non-significant values exhibit relatively high disease module sizes, with medians exceeding 100. However, the highest GDA score observed for non-significant values is 0.2, whereas significant disease modules still maintain large sizes when compared to the all-time significative scenario. We can highlight the case of malignant neoplasm of breast cancer, the only disease that exhibits a disease module at every GDA score threshold. This is notable, as maintaining a disease module above a threshold of 5 with such high GDA scores is rare and indicative of numerous associated genes. Initially, the module size is 6,941, but it is not considered significant until filtering at a GDA score of 0.3, which reduces the module size to 661. This example illustrates that a high module size alone does not guarantee significance (especially when derived from extensive literature) while a large module coupled with a GDA score above 0.3 is more likely to be meaningful.

In the last transition, “No → Yes → No”, all non-significant values begin at 0 but become significant at a GDA score of 0.1. Both cases exhibit a high disease module size, suggesting that this initial shift in significance could be attributed to the filtration of literature-based sources, which may not be the most reliable. The subsequent loss of significance is accompanied by a decrease in disease module size, reinforcing the idea that module size plays a key role in significance determination.

Both disease module size and GDA score are crucial factors in determining significance. In certain cases, a large disease module size compensates for a lower GDA score, while in others, a high GDA score counterbalances a smaller module size to maintain significance. Overall, most non-significant values cluster around a disease module size of 10, and very few non-significant values appear at a GDA score of 0.3 or higher. This suggests the emergence of a potential threshold, although further testing and analysis are required to establish a precise cutoff, which may also depend on the specific nature of each disease module.

## V. Conclusions

This study demonstrates that incorporating Gene-Disease Association (GDA) scores into the construction of disease modules significantly enhances the identification of robust and statistically significant sub-networks within the human interactome. The followed analysis indicates that applying a filtering threshold (specifically, retaining only those disease-associated genes with a GDA score above 0.3 and ensuring that the resulting modules contain more than 10 genes) substantially improves the overall significance of the disease modules.

The results reveal that modules meeting these criteria are less prone to the inclusion of only weak associations, particularly those derived predominantly from literature-based sources, favouring the inclusion of animal and curated sources. Moreover, the investigation into the relationship between module size and significance shows that larger modules tend to exhibit more stable statistical significance, suggesting that disease module size is a critical factor in module validation, especially if backed up by higher GDA scores.

In summary, the findings support the hypothesis that both the GDA score and module size are pivotal in determining the reliability of disease modules. These insights not only provide a framework for optimal seed gene prioritization but also have potential implications for enhancing target identification and drug repurposing strategies within network medicine.

## VI. Future steps and limitations

It is important to consider the limitations of this study when interpreting the obtained conclusions. First, the analysis is constrained by the inherent biases present in the underlying datasets. The interactome used in this study, derived from current PPI databases, is inevitably incomplete; many interactions, particularly those that are less studied or transient, may not be captured.

Future research should encompass extended validation of the proposed GDA score and module size thresholds across diverse diseases and datasets to evaluate their generalizability, as well as the integration of multi-omics data—such as transcriptomics, epigenomics, and metabolomics—to provide a more comprehensive view of disease mechanisms and improve gene prioritization. Additionally In this study, we constructed disease modules using the Largest Connected Component (LCC) approach, which is highly restrictive. Future research should explore alternative methods for defining disease modules to assess whether different approaches yield comparable or improved results.

## Acknowledgments

The work is a result of the project “Data-driven drug repositioning applying graph neural networks (3DR-GNN)” that is being developed under grant “PID2021-122659OB-I00” from the Spanish Ministerio de Ciencia e Innovación.

## Footnotes

1 https://disnet.ctb.upm.es/

2 https://medal.ctb.upm.es/internal/gitlab/disnet/network-medicine/network-medicine-and-disease-module-significance

